# A multiepitope fusion protein-based p-ELISA method for diagnosing bovine and goat brucellosis

**DOI:** 10.1101/2021.03.04.434020

**Authors:** Dehui Yin, Qiongqiong Bai, Xiling Wu, Han Li, Jihong Shao, Mingjun Sun, Jingpeng Zhang

## Abstract

In recent years, the incidence of brucellosis has increased annually, which has caused tremendous economic losses in agriculture and husbandry in various countries. Therefore, developing rapid, sensitive and specific diagnostic techniques for brucellosis has become critical brucellosis research. Bioinformatics technology was used to predict the B cell epitopes of the main antigen proteins of *Brucella*, and the validity of each epitope was verified by indirect enzyme-linked immunosorbent assay (iELISA). The verified epitopes were connected in series to construct a multiepitope fusion protein, goat, bovine brucellosis sera, and rabbit sera were collected to verify the antigenicity and specificity of this protein. Then, the fusion protein was used as a diagnostic antigen to construct paper-based ELISA (p-ELISA) technology. A total of 22 effective epitopes were predicted, and a fusion protein was successfully constructed, which showed good antigenicity and specificity. The constructed p-ELISA method was used for the simultaneous detection of bovine and goat brucellosis. ROC curve analysis showed that the sensitivity and specificity of protein detection in goat serum were 98.85% and 98.51%, respectively. The positive and the negative predictive value was 99.29% and 98.15%, respectively. When assessing bovine serum, the sensitivity and specificity were 97.85% and 96.61%, respectively. The positive and the negative predictive value was 98.28% and 97.33%, respectively. This study combined bioinformatics, fusion protein development and p-ELISA technologies to establish a sensitive and specific rapid diagnosis technology for brucellosis that can be used to assess the serum of bovine, goats and other livestock.

**IMPORTANCE:** Brucellosis has caused tremendous economic losses in agriculture and husbandry in various countries. Therefore, developing rapid, sensitive and specific diagnostic techniques for brucellosis has become critical brucellosis research. In this study, we used immunoinformatic technology to predict the B cell epitopes in the major outer membrane proteins of Brucella, synthesized polypeptides and coupled them with KLH, screened these polypeptides by iELISA methods, selected effective polypeptides as diagnostic antigens, and established a p-ELISA for brucellosis diagnosis based on a multiepitope fusion protein that can be used to assess the serum of bovine, goats and other livestock.

## 1 Introduction

Currently, a high incidence of brucellosis, a hazardous zoonosis, is reemerging worldwide, especially in developing countries, posing not only a large threat to human health but also tremendous losses in the world economy[1]. *Brucella* infection can be caused by humans directly contacting *Brucella*-infected goats, bovine, pigs and other livestock and their secretions or excreta or eating *Brucella*-contaminated food[2]. Due to the diversity of clinical manifestations of brucellosis and the lack of specific clinical manifestations, the diagnosis of brucellosis is very difficult, and it is easily misdiagnosed as other febrile diseases, such as dengue fever, malaria, or viral bleeding diseases[3,4].

There are many methods for diagnosing *Brucella*. Serological diagnostics are the most widely used and mature methods. However, serological diagnosis requires specific and sensitive antigens[5]. Currently, the commonly used antigens include whole-cell antigen and lipopolysaccharide (LPS). However, these antigens easily cross-react with the antibodies of other bacteria, which affects the specificity of the diagnosis. Therefore, it is very important to develop new diagnostic antigens to improve the specificity and sensitivity of serological diagnostic methods[6]. A large number of vaccine studies show that the *Brucella* outer membrane protein has good immunogenicity, which provides a direction for finding new diagnostic antigens[7–9]. The rapid development of paper-based enzyme-linked immunosorbent assay (p-ELISA) diagnostic methods based on paper chip technology provides a reference for the development of new brucellosis diagnostic methods[10].

In this study, five main antigen proteins of *Brucella* were screened, and the possible dominant B cell epitopes of these proteins were predicted by bioinformatics technology. We designed a new *Brucella* multiepitope fusion protein by concatenating the predicted dominant epitopes, making it a candidate antigen for the serological diagnosis of brucellosis. A rapid, sensitive and specific p-ELISA diagnostic technique for brucellosis that can detect protein in serum of bovine, goats and other livestock was successfully constructed.

## 2. Materials and methods

### 2.1 Serum samples

A total of 140 goat serum samples that were *Brucella* positive, 54 goat serum samples that were *Brucella* negative, 116 bovine serum samples that were *Brucella* positive, and 75 serum samples that were *Brucella* negative were provided by the China Animal Health and Epidemiology Center (Qingdao, China). All *Brucella*-positive and *Brucella*-negative sera were verified to be positive by the tube agglutination test and the Rose Bengal plate agglutination test (RBPT). All experiments involving animals or animal samples were fully compliant with ethical approval granted by the Animal Care and Ethics Committee of Xuzhou Medical University.

### 2.2 Prediction and synthesis of peptide epitopes

The *Brucella* outer membrane proteins (Omp) Omp16, Omp25, Omp31, Omp2b and BP26 were selected, and their amino acid sequences were obtained through the protein database at NCBI (https://www.ncbi.nlm.nih.gov/protein/). The conservation of the amino acid sequences was assessed by BLASTing. The prediction of B cell epitopes was carried out by using the B cell epitope prediction tool BepiPred linear epitope prediction 2.0 at IEDB (http://tools.iedb.org/bcell/). The predicted B cell epitope peptides were synthesized by Sangon Biotech (Shanghai, China) and coupled with keyhole limpet hemocyanin (KLH) with a purity of more than 90%.

### 2.3 Epitope antigenicity screening

Forty-five bovine and goat *Brucella*-positive sera were randomly selected to identify the antigenicity of the peptides through an indirect enzyme-linked immunosorbent assay (iELISA). In addition, a KLH carrier negative control and an LPS positive antigen control were established. For the procedure, in a 96-well microtiter plate (NUNC, Denmark), 100 μL of peptide (30 μg/mL in carbonate buffer solution (CBS), pH 9.6) was added to each well and incubated overnight at 4°C. The wells were blocked with 300 μL/well of 5% skimmed milk powder (Sangon, Shanghai) at 37°C for 2 hours, and then, 100 μL/well of *Brucella* serum was added (1:400 dilution with PBS) and incubated at 37°C for 1 hour. HRP-labeled protein G (diluted 1:5000, PBS) (Thermo, USA) was then added and incubated at room temperature for 30 min, after which an EL-TMB kit was utilized (Sangon). Optical density was measured at 450 nm (OD450) using an ELISA plate reader (BioTek, USA). After each step, the plates were washed 3 times with PBST.

### 2.4 Preparation of the fusion protein

The effective peptides were selected to be connected in series, and adjacent peptides were linked by the ‘GGGS’ linker. For the concatenated amino acid sequence, the molecular weight (https://web.expasy.org/compute_pi/), spatial conformation (http://zhanglab.ccmb.med.umich.edu/I-TASSER/) and other parameters were predicted. According to the amino acid sequence after concatenation, the codon was reversed, and the prokaryotic expression was optimized. The plasmid was constructed by whole gene synthesis and subcloned into expression vector pET30a (Beijing Protein Innovation, Beijing) and then transferred into competent cells (BL21 cells) for IPTG-induced expression. Specifically, competent cells (BL21 cells) (100 μL), stored at −80°C, were slowly thawed on ice, after which the ligation product was added to the cells and mixed well; the cells were then placed on ice for 30 min, heat shocked at 42°C for 90 s and then incubated in an ice bath for 2 min. Subsequently, 800 μL of nonresistant LB medium was added, incubated at 37°C for 45 min and centrifuged at 5000 rpm for 3 min. The majority of the supernatant was discarded, leaving approximately 100-150 μL, which was used to resuspended the cell pellet. The resuspended cells were added to LB plates with the corresponding resistance antibiotic and spread over plates, which were air-dried and cultured upside down and placed in an incubator at 37°C overnight. Then, the transformed BL21 cells were selected and cultured in 1.5 mL of LB liquid medium at 37°C and shaken at 200 rpm. The cells were incubated until the OD600=0.6, at which time they were induced by IPTG (0.5 mM) and cultured for 2 hours at 37°C. One milliliter of induced bacterial solution was centrifuged at 12000 rpm for 1 min, the supernatant was discarded, and the precipitate was resuspended in 50-100 μL of 10 mM Tris-HCl (pH 8.0) solution (the amount of added buffer was dependent on the amount of bacteria). Loading buffer equal to twice the volume of the resuspended precipitate was added, after which the sample was boiled at 100°C for 5 min and then assessed by SDS-PAGE electrophoresis.

After validation, 2 μL of activated bacterial solution was transferred to 750 mL of LB liquid medium at 37°C, spun at 200 rpm and incubated until the absorbance reached OD600 = 0.6-0.8. IPTG (0.5 mM) was added for overnight induction at 16°C. After centrifugation at 6000 rpm for 5 min, the supernatant was discarded, and the bacteria were collected. The bacteria were resuspended with 20-30 mL of 10 mM Tris-HCl (pH 8.0) solution and fragmented by ultrasonication (500 W, 60 times, 10 s each time, 15 s intervals). One hundred microliters of bacterial suspension was collected after ultrasonic treatment and centrifuged at 12000 rpm for 10 min. Fifty microliters of supernatant was poured into another EP tube. After the supernatant was removed, the precipitate was resuspended with 50 μL of 10 mM Tris-HCl (pH 8.0) solution and assessed by SDS-PAGE electrophoresis.

### 2.5 Purification of fusion protein

A nickel column (Ni Sepharose 6 Fast Flow, GE Healthcare) was washed with deionized water at pH 7.0. The nickel column was adjusted to pH 2~3. The column was washed with deionized water at pH 7.0. The nickel column was equilibrated with 10 mM Tris-HCl (pH 8.0) solution (approximately 100 mL). Then, the nickel column was equilibrated with a 10 mM Tris-HCl (pH 8.0) solution containing 0.5 M sodium chloride, (approximately 50 mL). Diluted sample was loaded. The sample contained sodium chloride at a final concentration of 0.5 M. After loading, the column was washed with 10 mM Tris-HCl (pH 8.0) solution containing 0.5 M sodium chloride. The proteins were eluted with a 10 mM Tris-HCl (pH 8.0) (containing 0.5 M sodium chloride) solution containing 15 mM imidazole, 60 mM imidazole, and 300 mM imidazole, and the protein peaks were collected separately. SDS-PAGE electrophoresis was used to assess the effect of protein purification.

### 2.6 Antigenicity assessment of the fusion protein

The iELISA method was used to assess the antigenicity of the purified protein. For the procedure, in a 96-well ELISA plate (NUNC, Denmark), 100 μL of fusion protein (2.5 μg/mL, CBS) was added to each well, and 100 μL of LPS (1 μg/mL, CBS) was added to the wells with positive antigen control and incubated overnight at 4°C. For blocking, 300 μL of 5% skimmed milk (PBS) was added per well and incubated at 37°C for 2 hours. Then, 100 μL of *Brucella*-positive serum (1:400 dilution, PBS) was added and incubated at 37°C for 1 hour; next, 100 μL of HRP-labeled protein G (diluted 1:8000, PBS) was added and incubated at room temperature for 30 min. After color development with the EL-TMB color kit, the absorbance of the wells was measured at OD450. After each step, the plates were washed 3 times with PBST.

At the same time, we used rabbit sera containing other pathogenic bacteria that easily cross-react with *Brucella*, including *Yersinia enterocolitica* O9, *Escherichia coli* O157:H7, *Salmonella*, *Vibrio cholerae*, *Vibrio parahaemolyticus*, and *Listeria monocytogenes*. All rabbit sera samples were purchased from Tianjin Biochip Corporation (Tianjin, China). The specificity of the protein was verified by iELISA. A 1:10000 dilution of HRP-labeled goat anti-rabbit secondary antibody (Bioworld, USA) was used in this assay, and the remainder of the steps were the same as outlined above.

### 2.7 Establishment of the p-ELISA method

A puncher was used to make a round sheet of Whatman No. 1 filter paper with a diameter of 10 mm, and a small hole (a diameter of 6 mm) was punched out of A4 plastic packaging paper. The 10 mm filter paper was placed in the center of the 6 mm hole in the plastic packaging paper, and a laminating machine joined the filter sheet and packaging paper, fixing and cutting the combined papers it into small strips with 3 holes in each strip. The fellow steps have been described in the literature[11] Five microliters of chitosan deionized water solution (0.25 mg/mL) was added to the round holes with Whatman No. 1 filter paper and dried at room temperature; then, 5 μL of 2.5% glutaraldehyde solution (PBS) was added, incubated at room temperature for 2 hours, and then washed 3 times with 20 μL of deionized water. Then, 5 μL of fusion protein solution (2.5 μg/mL, PBS) was added to each well, incubated at room temperature for 30 min, and washed 3 times with 20 μL of deionized water. Next, 20 μL of 5% skimmed milk powder was added and incubated for blocking at room temperature. Subsequently, 5 μL of serum (1:400 dilution) was added; after washing 3 times with PBST, 5 μL of HRP-labeled protein G (1:8000 dilution, PBS) was added, incubated at room temperature for 210 s, and washed 3 times with PBST. Finally, 5 μL of TMB substrate solution was added, and after 10 min, an HP Laser Jet Pro MFP M227 was used to scan the samples to obtain images. ImageJ software was used to perform gray intensity analysis for quantitation. The collected bovine and goat serum samples were assessed according to the established p-ELISA method, and ROC curves were used to analyze the diagnostic effect of the established method, including sensitivity and specificity.

### 2.8 Statistical analysis

Dot plot and receiver operating characteristic (ROC) analyses were performed using GraphPad Prism version 6.05 for Windows. The significance of gray intensity differences was determined by Student’s t-test (unpaired t-test). Differences were considered statistically significant when P< 0.05.

## 3 Results

### 3.1 B cell epitope peptide prediction and antigenicity verification

A total of 22 B cell epitopes were predicted, including BP26, Omp16, Omp25, Omp31, and Omp2b, which were predicted to have 6, 2, 5, 5, and 4 epitopes (table S1), respectively. Indirect ELISA results showed that all 22 peptides recognized portions of the serum (Fig. 1)

**Fig. 1.**
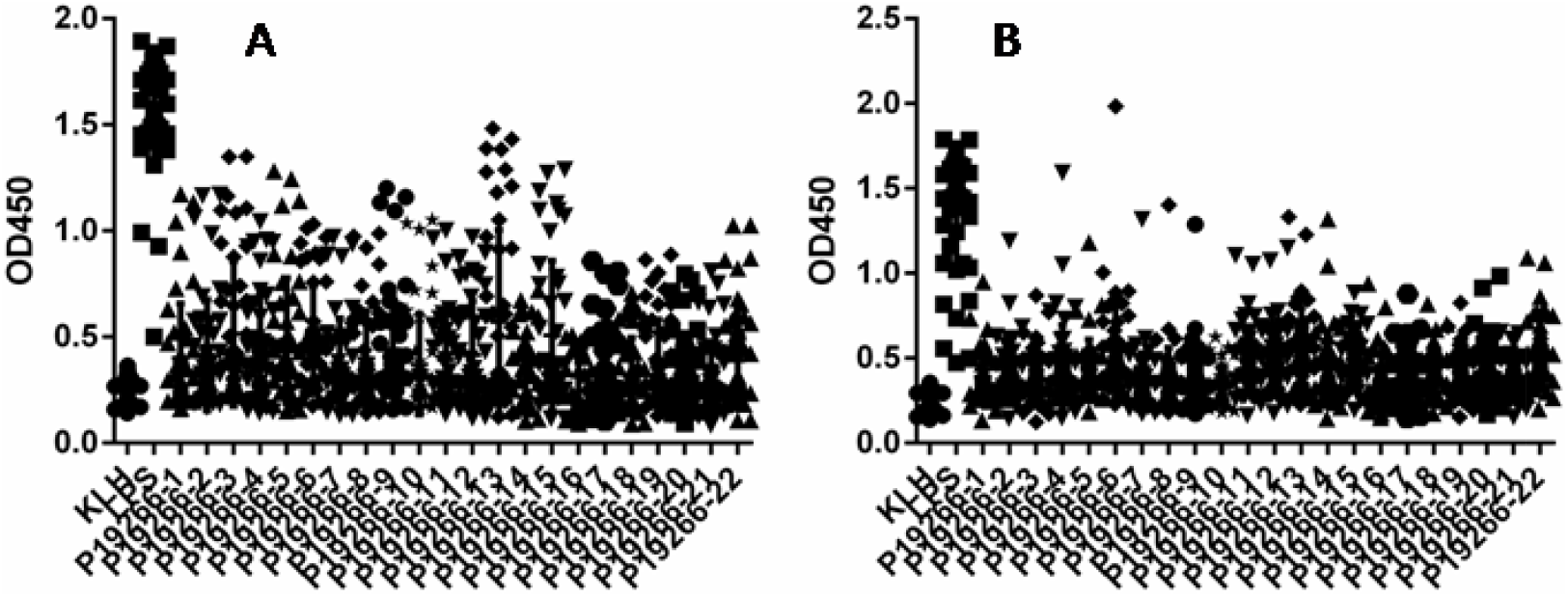
The results of iELISA of each peptide identification-positive brucellosis serum. (A) Sheep brucellosis serum. (B) Bovine brucellosis serum.

### 3.2 Preparation of the multiepitope fusion protein

The 22 epitopes were connected in series, and the adjacent peptides were connected with the ‘GGGS’ linker to form an amino acid sequence of the fusion protein (Figure S1). Prokaryotic expression was induced by IPTG, and the target protein was expressed in the supernatant. SDS-PAGE (15%) electrophoresis showed a protein band with an approximate molecular weight of 66 kD. After mass spectrometry analysis, it was confirmed that this band was the target protein. After purification, most of the miscellaneous bands were removed, and gray intensity analysis showed that the purity of the purified protein was approximately 90% (Fig. 2).

**Fig. 2.**
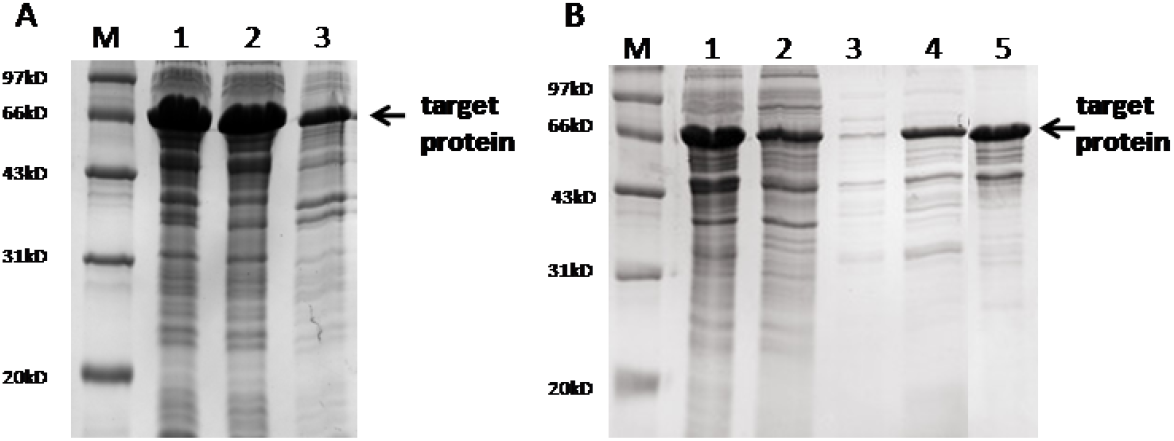
SDS-PAGE analysis of fusion protein. (A) Protein expression results. M, marker; lane 1, whole bacteria after ultrasound; lane 2, supernatant after ultrasound; lane 3, precipitation after ultrasound. (B) SDS-PAGE after protein purification. M, marker; lane 1, the original protein before purification; lane 2, flow-through solution; lane 3,15mM imidazole elution fraction; lane 4,60mM imidazole elution fraction; lane 5,300mM imidazole elution fraction.

### 3.3 Antigenicity assessment of the fusion protein

After iELISA analysis, when the fusion protein was used as an antigen to test its diagnostic ability with goat serum, the area under the ROC curve was 0.9799 (95% CI, 0.9654 to 0.9944), and the cutoff value calculated by the Youden Index was 0.4675. In this case, the diagnostic sensitivity was 87.14% (95% CI, 0.8044 to 0.9220), and the specificity was 100.0% (95% CI, 0.9340 to 1.000). The positive predictive value as 100.0%, and the negative predictive value was 75.00%. When LPS was used as an antigen to test its diagnostic ability with goat serum, the area under the ROC curve was 0.9514 (95% CI, 0.9191 to 0.9836), the cutoff value was 0.8890, the diagnostic sensitivity was 82.00% (95% CI, 0.7305 to 0.8897), and the specificity was 95.83% (95% CI, 0.8575 to 0.9949). The positive predictive value was 98.39%, and the negative predictive value was 74.29% (Fig. 3 and table 1).

**Fig. 3.**
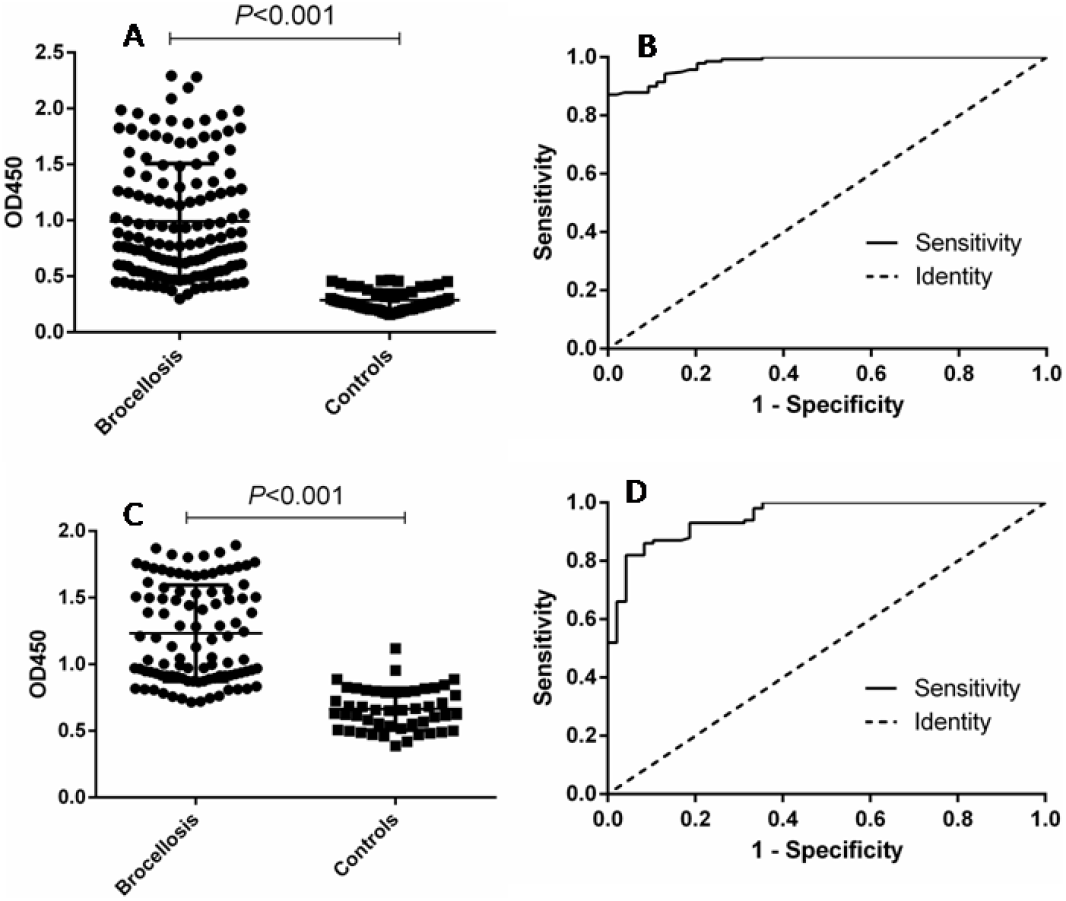
ELISA analysis of goat serum samples. (A) Dotplot of the fusion protein ELISA assay. (B) ROC analysis of fusion protein IELISA assay results. (**C)** Dotplot of the LPS antigen ELISA assay. (D) ROC analysis of LPS antigen ELISA assay results.

**Table 1.** Positive and negative predictive values of the test calculated for different cut-off values

| Cut-off value | Positive |  | Negative |  | PPV (%) | NPV (%) |
| --- | --- | --- | --- | --- | --- | --- |
|  | TP | FN | TN | FP |  |  |
| $\geq 0.4675$(Fusion protein)<sup>a</sup> | 122 | 18 | 54 | 0 | 100.0 | 75.00 |
| $\geq 0.8890$(LPS)<sup>a</sup> | 122 | 18 | 52 | 2 | 98.39 | 74.29 |
| $\geq 0.4530$(Fusion protein)<sup>b</sup> | 103 | 13 | 70 | 5 | 95.37 | 84.34 |
| $\geq 0.8105$(LPS)<sup>b</sup> | 107 | 9 | 68 | 7 | 93.86 | 88.31 |
| $\geq 34.12$(p-ELISA)<sup>a</sup> | 139 | 1 | 53 | 1 | 99.29 | 98.15 |
| $\geq 30.21$(p-ELISA)<sup>b</sup> | 114 | 2 | 73 | 2 | 98.28 | 97.33 |

a, goat sera; b, cattle sera. TP, true positives; TN, true negatives; FP, false positives; FN, false negatives; PPV, positive predictive value $(TP/TP+FP) \times 100$ ; NPV, negative predictive value $(TN/TN+FN) \times 100$ .

To test its diagnostic ability with bovine serum, the area under the ROC curve was 0.9518 (95% CI, 0.9224 to 0.9812), and the cutoff value calculated by the Youden index was 0.4530. In this case, the diagnostic sensitivity was 88.79% (95% CI, 0.8160 to 0.9390), and the specificity was 93.33% (95% CI, 0.8512 to 0.9780). The positive predictive value was 95.37%, and the negative predictive value was 84.34%. When LPS was used as an antigen to diagnose bovine serum, the area under the ROC curve was 0.9528 (95% CI, 0.9187 to 0.9868), and the cutoff value was 0.8105. In this case, the diagnostic sensitivity was 90.63% (95% CI, 0.8295 to 0.9562), and the specificity was 90.28% (95% CI, 0.8099 to 0.9600). The positive predictive value was 93.86%, and the negative predictive value was 88.31% (Fig. 4 and table 1).

**Fig 4.**
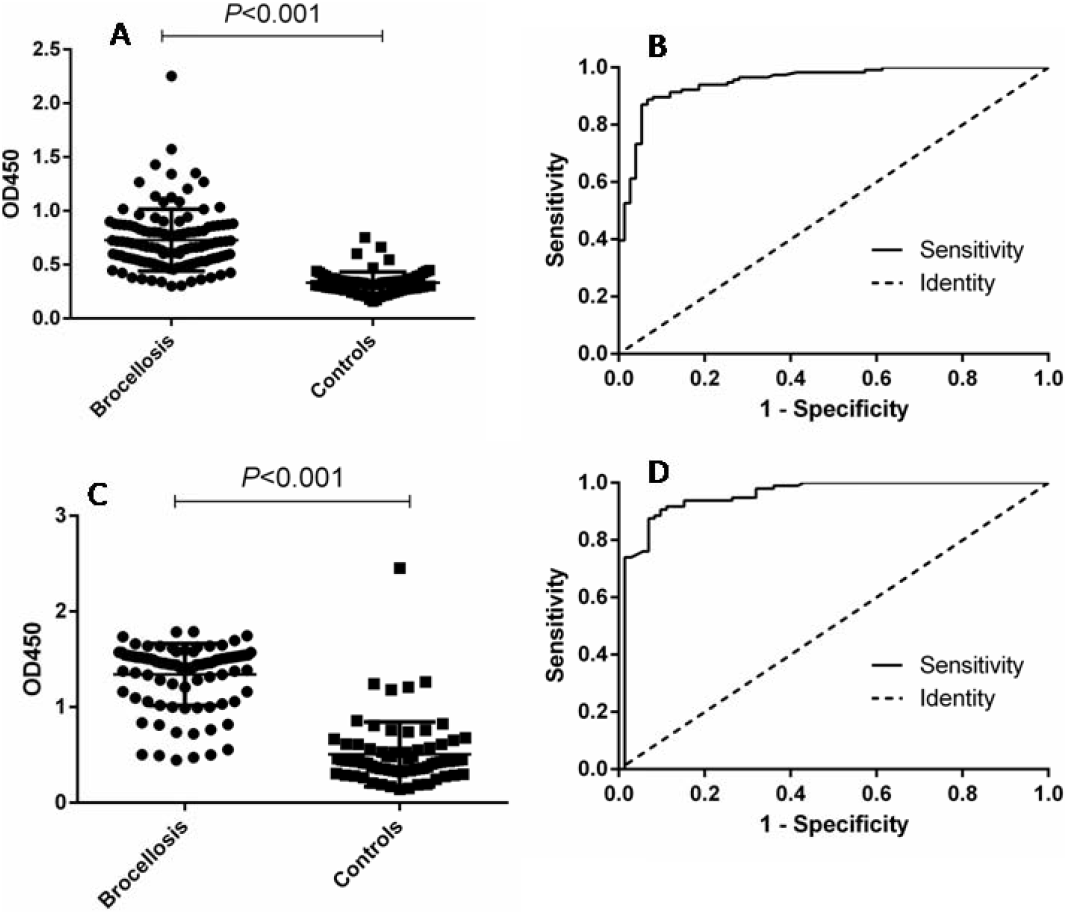
ELISA analysis of cattle serum samples. (A) Dotplot of the fusion protein ELISA assay. (B) ROC analysis of fusion protein IELISA assay results. (**C)** Dotplot of the LPS antigen ELISA assay. (D) ROC analysis of LPS antigen ELISA assay results.

### 3.4 Determining the cross-reactivity with the fusion protein

To verify whether the fusion protein as a diagnostic antigen shows cross-reactivity with other bacteria, we selected 6 zoonotic pathogens for a cross-reactivity test. The results showed that the fusion protein did not cross-react with other bacteria according to an S/N (OD450, sample/negative) > 2.1, which indicated that the fusion protein as a diagnostic antigen has good specificity (table 2).

**Table 2.** Specific cross-reactivity test results of the indirect ELISA diagnostic method for the fusion protein.

| Rabbit sample | OD450 | S/N |
| --- | --- | --- |
| *Vibrio parahaemolyticus* | 0.1230 | 1.64 |
| *Escherichia coli* O157:H7 | 0.0457 | 0.61 |
| *Salmonella* | 0.1267 | 1.69 |
| *Vibrio cholerae* | 0.0598 | 0.80 |
| *Yersinia enterocolitica* O9 | 0.0443 | 0.59 |
| *Listeria monocytogenes* | 0.0758 | 1.01 |
| Negative | 0.0751 | -- |

### 3.5 Evaluation of the diagnostic ability of the p-ELISA

To verify the effectiveness of the established p-ELISA method, the collected serum samples were tested. When diagnosing goat sera, the area under the ROC curve was 0.9986 (95% CI, 0.9957 to 1.002). The cutoff value was 34.12, the diagnostic sensitivity was 98.85% (95% CI, 0.9376 to 0.9997), and the specificity was 98.51% (95% CI, 0.9196 to 0.9996). The positive predictive value was 99.29%, and the negative predictive value was 98.15% (table 1). When testing its activity in bovine serum, the area under the ROC curve was 0.9964 (95% CI, 0.9910 to 1.002), and the cutoff value calculated by the Youden index was 30.21. In this case, the diagnostic sensitivity was 97.85% (95% CI, 0.9245 to 1.002), and the specificity was 96.61% (95% CI, 0.8829 to 0.9959). The positive predictive value was 98.28%, and the negative predictive value was 97.33% (table 1).

## 4 Discussion

Brucellosis is a serious zoonotic disease. Bovine and goats are the most commonly infected animals[12]. Currently, culling infected livestock is widely used to eradicate brucellosis worldwide[13]. The accurate diagnosis of livestock is essential for the prevention and control of the disease and for reducing unnecessary economic losses. Particularly in China, where the base of bovine and goats is relatively large, fast and efficient screening methods are of great significance[14]. Serological diagnostic techniques, mainly the agglutination test, RBPT, complement fixation test (CFT), ELISA, immunochromatographic diagnostic test (ICDT), and fluorescence polarization assay (FPA), are currently the commonly used screening methods for brucellosis[15,16]. Serological methods have the advantages of high sensitivity and a short operation time. Serological methods are the most commonly used methods for diagnosing brucellosis, but the existing serological diagnostic methods have the shortcomings of cross-reactivity with other bacteria (such as *Escherichia coli* O157:H7 and *Yersinia enterocolitica* O9)[17], which easily results in false positives. Reducing cross-reactions with other bacterial species is the key to improving the diagnostic specificity of serological methods.

ELISA is currently the most widely studied serological diagnosis method, even as diagnostic confirmation in brucellosis[16]. The main problem with using ELISAs for the diagnosis of brucellosis is the choice of antigen, but to date, ELISA-based diagnoses lack a single standard antigen[18]. Currently, the most commonly used diagnostic antigens used in ELISA are whole bacteria or extracts. These diagnostic antigens are prone to cross-reactivity with other bacteria, have poor specificity and have considerable defects. Therefore, the development of new diagnostic antigens is key to improving the diagnostic effect of ELISAs.

B cell epitopes refer to antigen regions that can be recognized by B cell receptors (BCRs) or antibodies produced by the humoral immune system. Epitope recognition is an important aspect of immunology. The rapid development of bioinformatics provides convenient ways and methods for epitope prediction[19]. Bioinformatics tools can be used to select potential cell epitopes without cultivating pathogens, which has the advantages of rapid testing and low cost[20]. Currently, B cell epitope prediction tools include databases, algorithms, and web servers. In this study, five main antigen components of *Brucella* were selected, and 22 effective epitopes were successfully predicted using the IEDB website. After iELISA verification, each peptide was used to identify a certain sample of brucellosis serum. Therefore, combining these peptides to construct new protein antigens can theoretically increase the recognition of proteins in serum. The results of the iELISA confirmed our hypothesis. The diagnostic specificity of this protein was higher than that of LPS, and it did not cross-react with other bacteria. The fusion protein we constructed has potential value as a diagnostic antigen.

In recent years, new disease diagnosis and pathogen detection technologies have been developed based on ELISA. Among these technologies, the p-ELISA has been rapidly developed and widely used in related fields such as health testing and diagnosis[21]. The p-ELISA method is a new technology developed based on the traditional ELISA method principle, using paper as the solid-phase carrier[22]. Compared with the traditional ELISA method, p-ELISA is faster and can be used to detect proteins within 1 hour. Less reagent and only 2-5 μL of the samples are required, no special instruments such as microplate readers are needed (only smart phones or cameras are needed to take photos, combined with Photoshop and other software to complete the data analysis), and the carrier is replaced with harmless easy-to-handle paper, which greatly saves costs and provides a new research direction for the development of new methods for the diagnosis of brucellosis [23]. Currently, the most common paper-processing method of the p-ELISA method involves constructing hydrophilic and hydrophobic areas through wax printing technology. This method often requires expensive printers, which limits the application of this method. We use plastic-encapsulated paper to construct the hydrophobic area, punching small holes into it, filling the small holes with hydrophilic paper sheets to make a sandwich structure. The paper can be processed with an ordinary laminator; therefore, the cost is low. Through serum detection, p-ELISA has a better diagnostic outcome. Compared with the traditional ELISA method, it has improved sensitivity and specificity, and the dosage of the reagents and the time of the operation are significantly reduced. This method can simultaneously detect bovine and goat brucellosis and has certain application prospects.

In conclusion, we predicted 22 B cell epitopes by using bioinformatics technology and successfully constructed a *Brucella* diagnostic antigen. Combined with this antigen, we constructed a p-ELISA method that can be used to diagnose bovine and goat brucellosis simultaneously. However, currently, it is extremely difficult to identify all affected animals from epidemiological units through serological tests, and there is no serological test able to differentiate between affected and vaccinated animals. The p-ELISA method we have constructed cannot yet solve the two questions mentioned above. In addition, our study is only preliminary, the number of samples is small, and the selected samples, especially *Brucella*-positive samples, represent highly positive samples. The effectiveness of this method needs further study, such as by testing a large number of random samples.

## Conflict of Interest

The authors declare no potential conflicts of interest with respect to the research, authorship, and/or publication of this article.

## Funding

This work was supported by the National Natural Science Foundation of China (Grant number 81802101). The funders had no role in the study design, data collection and analysis, decision to publish, or preparation of the manuscript.

